# Event-Related Phase Synchronization Propagates Rapidly across Human Ventral Visual Cortex

**DOI:** 10.1101/2021.08.02.454804

**Authors:** Oscar Woolnough, Kiefer J. Forseth, Patrick S. Rollo, Zachary J. Roccaforte, Nitin Tandon

## Abstract

Visual inputs to early visual cortex integrate with semantic, linguistic and memory inputs in higher visual cortex, in a manner that is rapid and accurate, and enables complex computations such as face recognition and word reading. This implies the existence of fundamental organizational principles that enable such efficiency. To elaborate on this, we performed intracranial recordings in 82 individuals while they performed tasks of varying visual and cognitive complexity. We discovered that visual inputs induce highly organized posterior-to-anterior propagating patterns of phase modulation across the ventral occipitotemporal cortex. At individual electrodes there was a stereotyped temporal pattern of phase progression following both stimulus onset and offset, consistent across trials and tasks. The phase of low frequency activity in anterior regions was predicted by the prior phase in posterior cortical regions. This spatiotemporal propagation of phase likely serves as a feed-forward organizational influence enabling the integration of information across the ventral visual stream. This phase modulation manifests as the early components of the event related potential; one of the most commonly used measures in human electrophysiology. These findings illuminate fundamental organizational principles of the higher order visual system that enable the rapid recognition and characterization of a variety of inputs.

## Introduction

Ventral occipitotemporal cortex is organized in a cortical hierarchy from early visual regions (e.g. calcarine cortex) to associative areas (e.g. fusiform cortex) (Felleman and Van Essen, 1991; Morán et al., 1987), that constitutes the ventral visual stream (Mishkin et al., 1983). This processing pathway is crucial for the integration of visual processing with language and memory (Forseth et al., 2018; Ghuman et al., 2014; Hirshorn et al., 2016; Kadipasaoglu et al., 2016; Tang et al., 2014; Woolnough et al., 2021b), but it is unclear how the propagation of information between functional components of the human ventral visual pathway is coordinated.

Recently, spatial propagation of information has been characterized, in both humans and non-human primates, as travelling waves of cortical activation. These occur at both the macro and micro scale and represent the structured propagation of information across the cortical surface in response to inputs (Lozano-Soldevilla and VanRullen, 2019; Muller et al., 2018, 2014; Sato et al., 2012) or may occur spontaneously (Bahramisharif et al., 2013; Davis et al., 2020; Halgren et al., 2019; Muller et al., 2016; Zhang et al., 2018). In the ventral visual stream of macaques this manifests as a rapid feedforward sweep of activation following visual stimulation (Lamme and Roelfsema, 2000). This spatiotemporal organization, manifest as a travelling wave indexes synchronized long-range cortico-cortical connections, enabling the integration of information across multiple cortical areas (Sato et al., 2012; Zhang et al., 2018) and perhaps also facilitates predictive coding (Alamia and VanRullen, 2019; Arnal and Giraud, 2012). Such travelling waves have commonly been characterized as a spatial propagation of phase. Sensory inputs result in the modulation of phase of ongoing oscillations (Howard and Poeppel, 2012; Lakatos et al., 2008, 2005; Luo et al., 2010) and the timing of inputs relative to the phase of ongoing oscillations impacts perception (Bonnefond and Jensen, 2012; Davis et al., 2020; Dugué et al., 2015; Forseth et al., 2020; Mercier et al., 2015). Based on this, we hypothesize there is a stimulus-evoked modulation of phase that spatiotemporally propagates through the human ventral visual pathway.

To probe the spread of visually-evoked phase modulation across the ventral visual stream, we utilized high spatiotemporal resolution intracranial recordings in a large human cohort (82 patients), with almost ubiquitous coverage of the ventral visual stream (1,929 electrodes), who performed a language task with relatively low visual complexity stimulus (word reading) or a memory task with a high complexity visual stimulus (face naming).

## Results

82 patients semi-chronically implanted with subdural grid electrodes (SDEs; 14 patients) or stereotactically placed depth electrodes (sEEGs; 68 patients) for the localization of intractable epilepsy performed word reading (n = 40) and face naming (n = 57) tasks (Figure 1A,B). Phase modulation was quantified by computing the inter-trial phase coherence (ITC; Figure 1E,F) of the broadband low frequency signal (2-30 Hz) at each electrode, across multiple trials.

**Figure 1:**
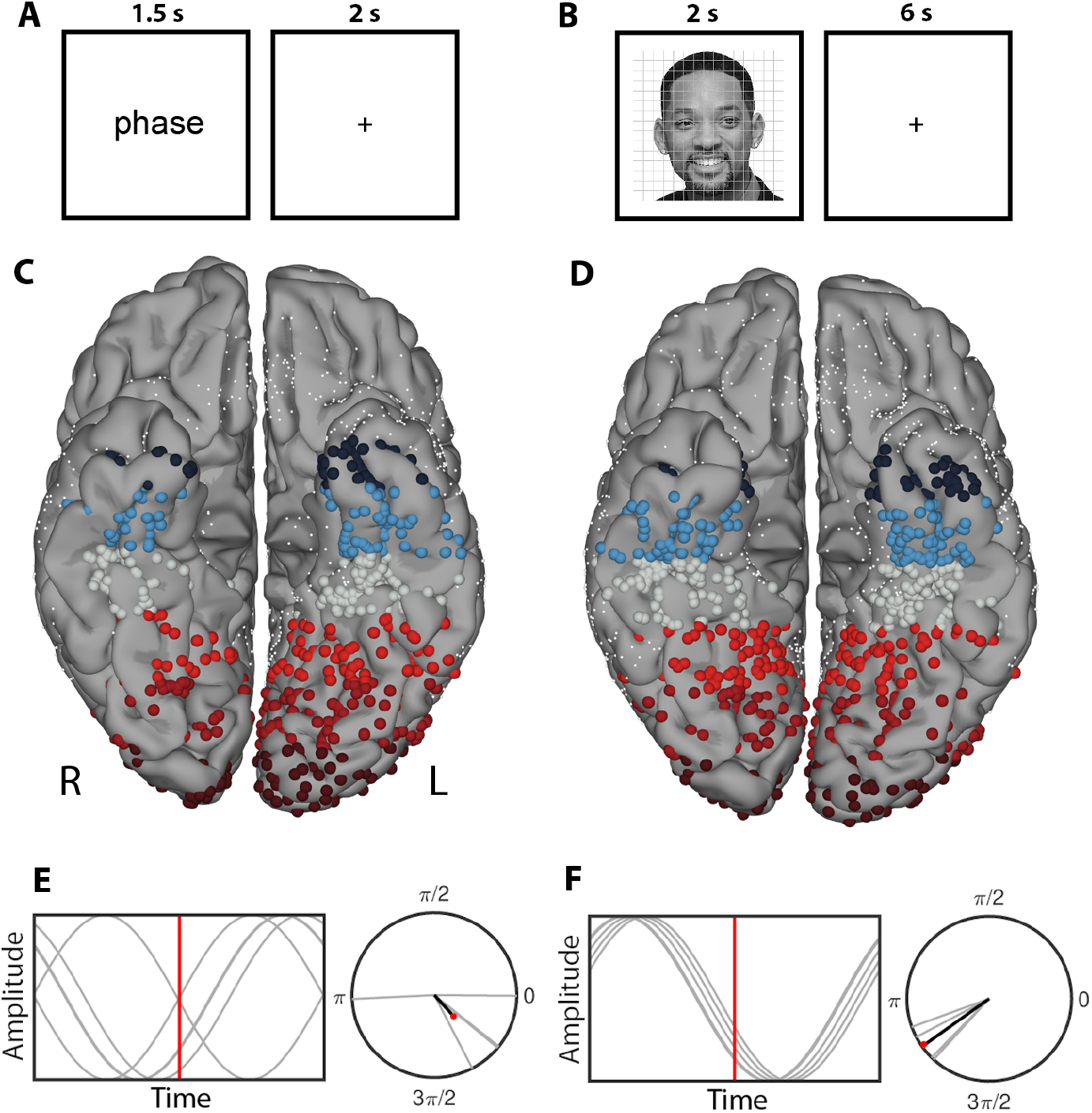
Tasks, Patients and ITC Analysis. (A,B) Schematic representation of the words (A) and faces (B) tasks. (C,D) Individual electrode locations, within the ventral ROI, for patients reading words (C; 1,059 electrodes, 40 patients) and recognizing faces (D; 1,246 electrodes, 57 patients) plotted on a standardized N27 brain. Electrode colors represent the relative locations with individual electrodes clustered into 20 mm bins along the y-axis. Smaller white spheres represent electrodes outside the ventral ROI. (E,F) Schematic representation with artificial data to illustrate low (E; ITC = 0.25) and high (F; ITC = 0.95) phase alignment along with a representation of the vector mean.

### Phase Modulation induced by Words and Faces

Data from two representative patients, with broad coverage of ventral occipitotemporal cortex, are shown in Figure 2. In both instances, increases in inter-trial phase coherence were seen in electrodes from occipital pole to mid-fusiform cortex, for both tasks. Within each patient we also observed highly comparable patterns of ITC across both tasks. As expected, during the period preceding the onset of visual stimulation, the trial-by-trial distribution of phases at any given time was reasonably uniform. However, following presentation of a visual stimulus, they tended to a coherent center frequency (Figure 2C,D). This phase coherence persisted for approximately 2 cycles (4π rads, ∼300ms) (Figure 2F,H). The temporal progression of the coherent center frequency was highly similar across both tasks.

**Figure 2:**
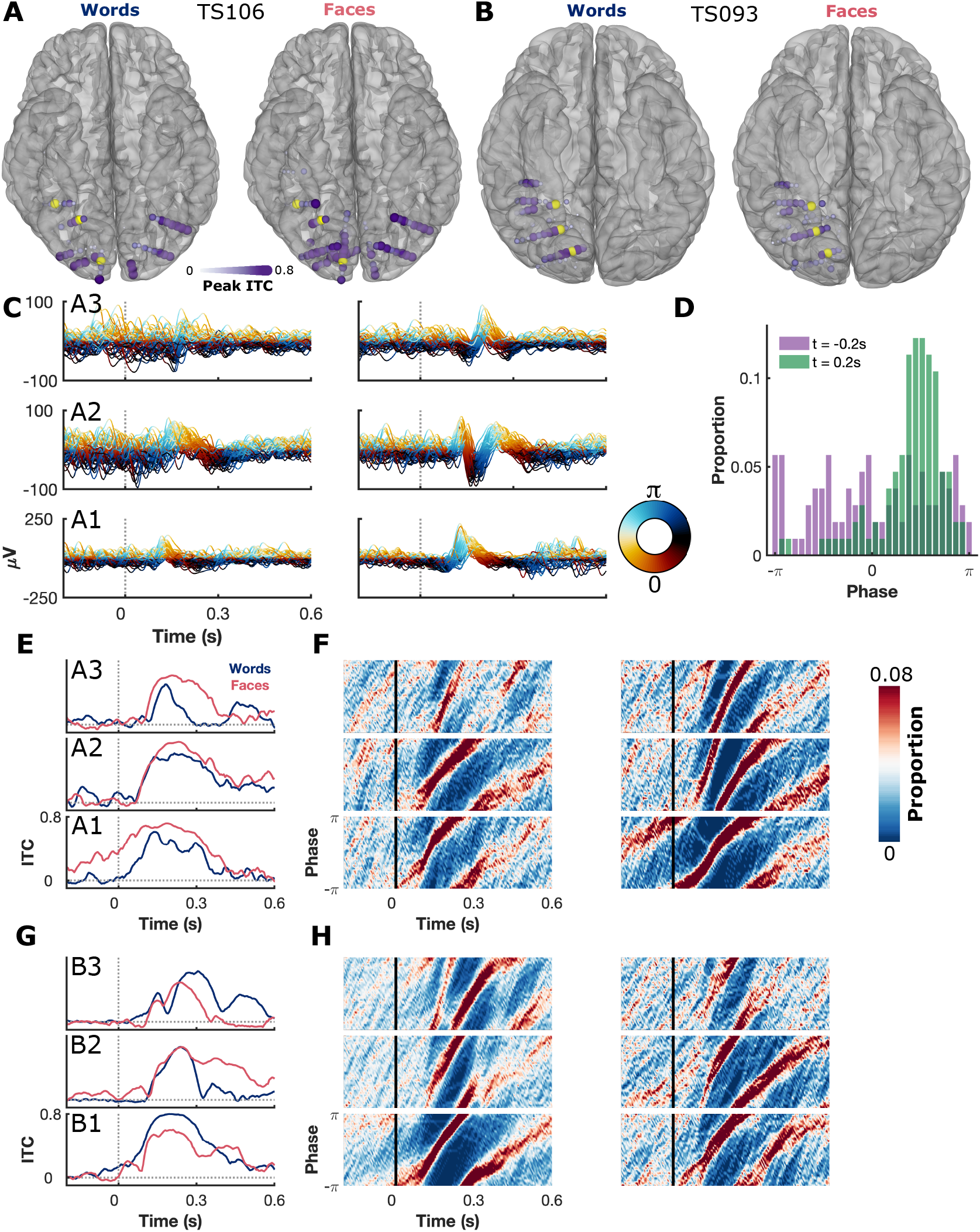
Spatial Propagation of Phase Coherence in Exemplar Patients. (A,B) ITC distributions, in native patient space, for TS106 (A) and TS093 (B). Results are shown for words (left) and faces (right). (C) Broadband low frequency (2-30 Hz) signals of 60 trials from TS106 from the words (left) and faces (right) tasks. Traces colored based on instantaneous generalized phase. (D) Phase distribution for a representative electrode at two time points during the faces task. (E-H) ITC (E,G) and phase distributions (F,H), with words (left) and faces (right) shown for three electrodes each from TS106 (E,F) and TS097 (G,H). Electrodes are highlighted in yellow in the brain plots and electrode numbers increase from posterior to anterior.

To evaluate the consistency of these effects across the population (n = 82 patients), we represented ITC amplitudes on a group normalized cortical surface from occipital pole to anterior fusiform cortex (Figure 3A,B). High ITC (>0.5) values were noted in associative cortex as far anterior as mid-fusiform. Stimulus presentation resulted in two distinct peaks of the ITC measure – a higher one around stimulus onset and a smaller one at offset. Differences in offset time between the two tasks, based on experimental design, confirmed that this was an offset response (Figure 3D,E). Results calculated using the raw signal (0.3-100Hz) showed highly correlated peak ITC with the broadband low frequency (2-30Hz) signal (Spearman correlation; words, r(1,059) = 0.86, p < 0.001; faces, r(1,246) = 0.82, p < 0.001; Supplementary Figure 1E,F) but with the raw signal displaying a longer duration of significant ITC (Supplementary Figure 1G,H).

**Figure 3:**
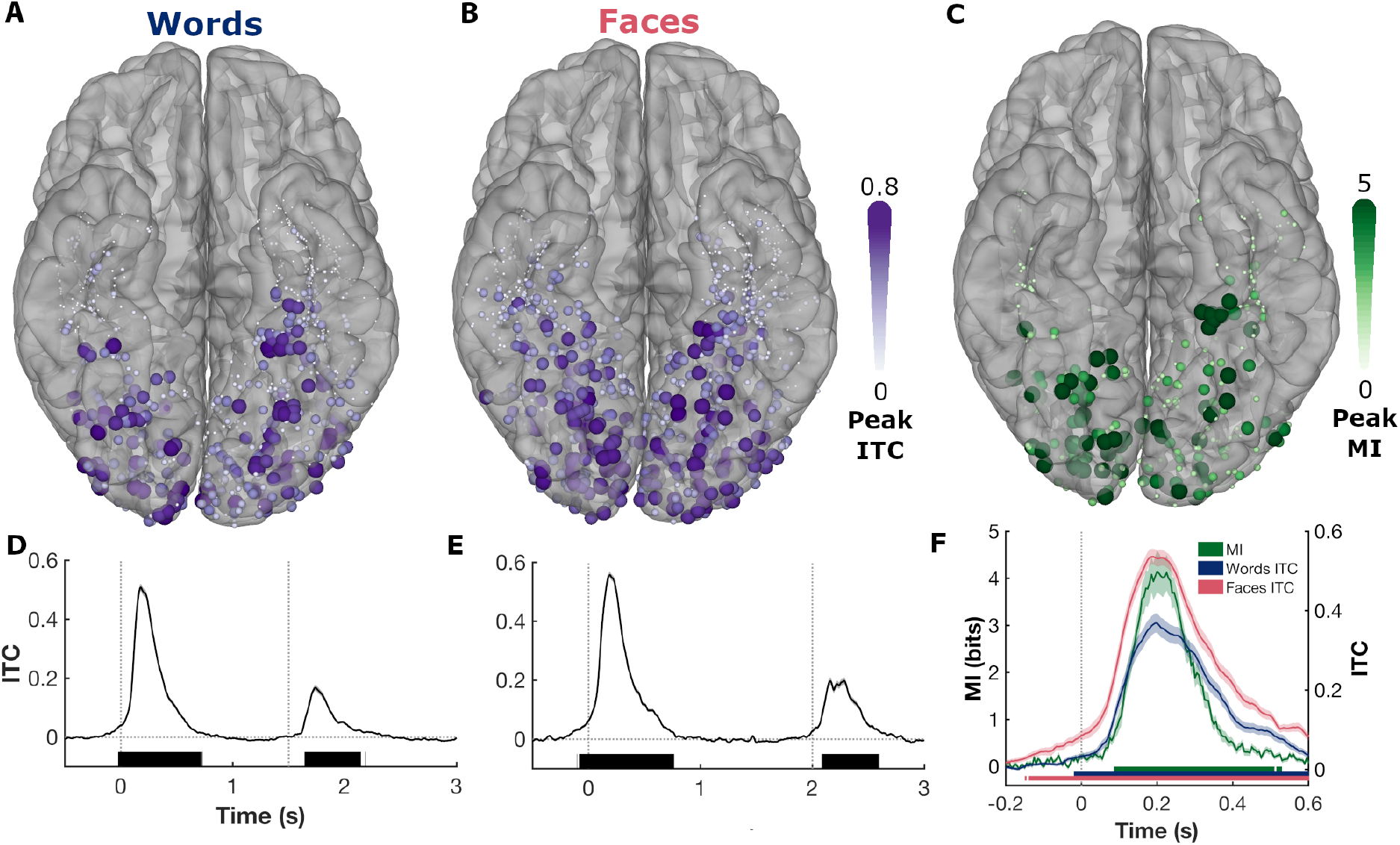
Population Phase Coherence. (A,B) Peak ITC at individual electrodes for words (A; 1,059 electrodes) and faces (B; 1,246 electrodes) tasks represented on a standardized surface. (C) Peak mutual information (MI) between the phase distributions in response to word or face presentation in patients who performed both tasks (354 electrodes, 15 patients). (D,E) Time course of ITC (mean ±SE) at electrodes with the top 20% peak amplitude ITC responses to words (D) and faces (E). Vertical lines denote the onset and offset times of each stimulus. Colored bars represent regions of significant increase over baseline (q<0.001). (F) MI and ITC for the electrodes with the top 20% highest peak MI. See also Supplementary Figures 1, 2.

The results thus far demonstrate that within each task there is phase coherence between trials and the magnitude and duration of this coherence is reasonably conserved across different tasks. But, how consistent is this pattern of evoked phase across tasks? To quantify this, we measured mutual information (MI) in patients who performed both tasks (n = 15). We observed an increase in MI following the visual onsets, representing an increase in the similarity of phase distributions between tasks. This inter-task coherence peaked at ∼200ms following visual onset, highly concordant with the within-trial ITC (Figure 3F). 9 out of the 15 patients had at least one electrode peaking at >5 bits (48/354 electrodes; Supplementary Figure 2A). Peak MI correlated well with peak ITC, measured across both tasks (Spearman correlation, r(354) = 0.76, p<0.001; Supplementary Figure 2C).

Next, we sought to characterize the spatiotemporal properties of this response, by dividing the ventral cortical surface into 20 mm bins along the y axis of the normalized surface in Talairach space, and used this to compare grouped ITC profiles from each bin (Figure 4). We noted a pronounced posterior-to-anterior gradient of both amplitude and onset latency of ITC with the largest responses in early visual cortex and progressive diminution anteriorly in both hemispheres, for both tasks. A 4D, cortical surface-based representation of the propagation of this phase coherence across the cortical surface revealed a posterior-to-anterior wave, highly conserved across both tasks (Movie 1).

**Figure 4:**
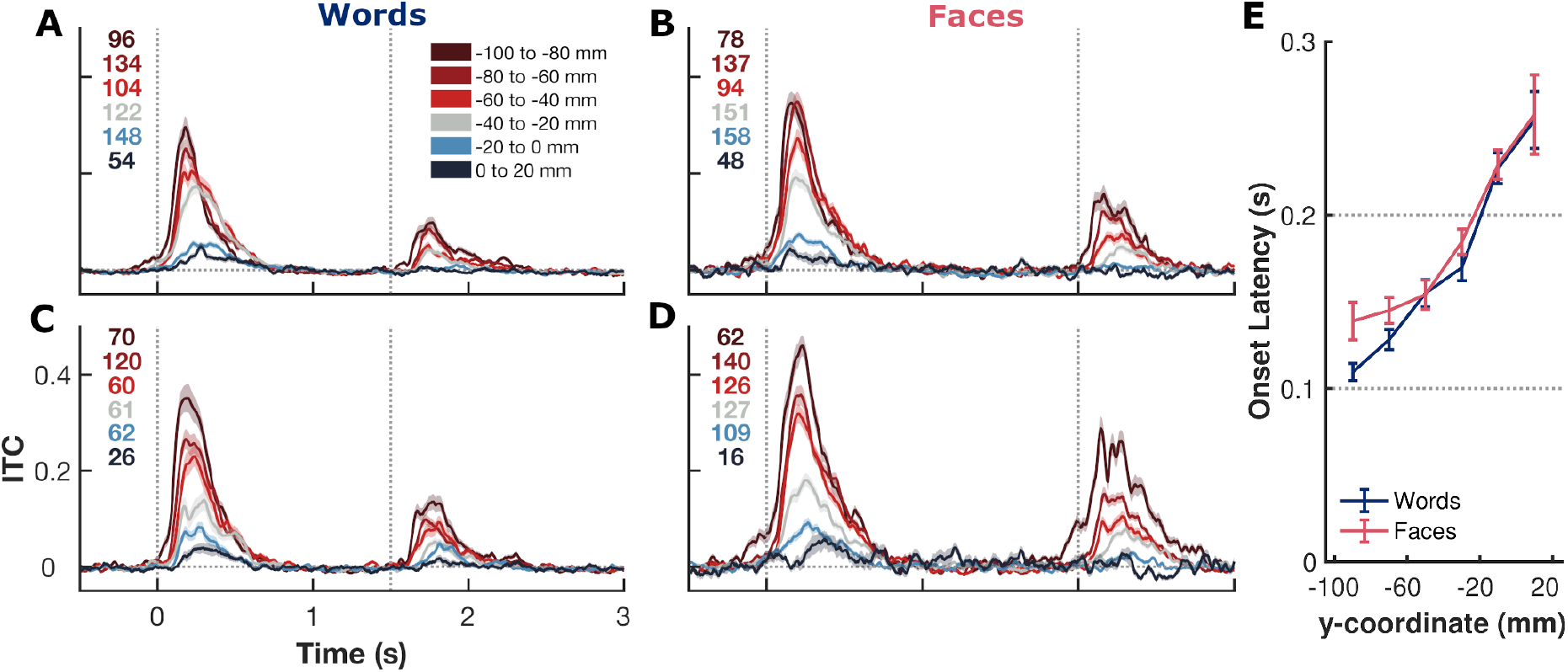
ITC – Positional Analysis. (A-D) ITC in response to words (A,C) or faces (B,D) as a function of anteroposterior position in the left (A,B) and right (C,D) hemisphere. Presented as mean ± SE. Number of electrodes in each ROI shown. Vertical lines denote stimulus onset and offset times. (E) ITC demonstrates a posterior-to-anterior gradient of first onset latency (ITC first-derivative >3.5SD deviation from baseline).

Movie 1: **Spread of Phase Coherence across the Cortical Surface**. Population-level, surface-based representation of ITC in response to words (left) and faces (right).

To quantify this posterior-to-anterior latency gradient, we calculated the time point at which the first derivative of the ITC first significantly deviated from baseline at each electrode (Figure 4E). In both tasks there was a clear gradient from calcarine to anterior fusiform. Onset latency showed a significant association with position in the anteroposterior axis for both words (t(781) = 12.5, β = 1.0 ± 0.08, p < 0.001, r^2^ = 0.30) and faces (t(779) = 9.7, β = 1.3 ± 0.13, p < 0.001, r^2^ = 0.21) tasks. This reflects a posterior-to-anterior progression of the timing of ITC across the cortical surface from early sensory cortex to associative regions and a consistent propagation rate of approximately 0.8-1 m/s in Euclidean space.

To further investigate the spatiotemporal consistency of this effect between tasks, we analyzed just the subset of patients who performed both tasks (n = 15). There was no significant difference in ITC onset latency between the two tasks (t(462) = −1.6, β = −0.03 ± 0.02, p = 0.12) or any significant effect of task on the propagation rate (t(462) = −1.3, β = −0.35 ± 0.27, p = 0.20).

To further characterize this posterior-to-anterior spread and to identify the degree of propagation of phase information across the cortical surface, we calculated the lagged phase locking value (PLV) between pairs of electrodes. By lagging the time courses of phase progression between pairs of electrodes we sought to determine to what degree the future phase of anterior sites could be predicted by the current phase of posterior sites (Figure 5A). This analysis demonstrated a strong bias toward a posterior-to-anterior spread of phase information, with the phase of anterior sites being driven by more posterior regions (Figure 5B, Supplementary Figure 3).

**Figure 5:**
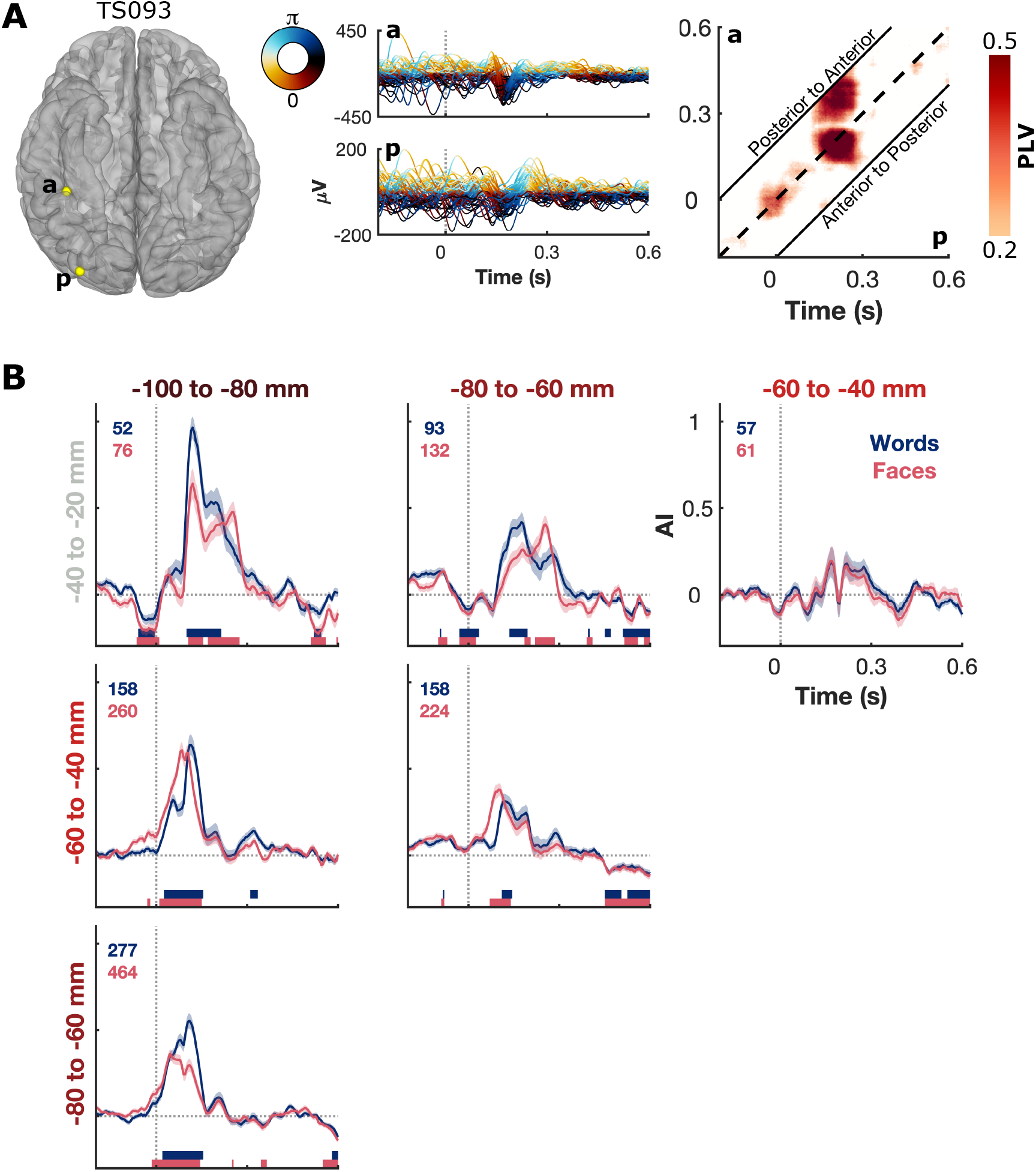
Propagation of Phase. (A) Exemplar electrode pair demonstrating the lagged phase locking value (PLV) analysis. PLV was calculated between the instantaneous phase of the posterior electrode (p) and the time lagged phase of the anterior electrode (a; −200 to 200 ms). Greater PLV above the center diagonal (dashed line) represents a greater ability to predict future phase of the anterior electrode based on the current phase of the posterior electrode. (B) Calculated asymmetry index (AI) of PLV above and below the center diagonal for high ITC (>0.5) electrode pairs across ROIs. AI > 0 represents a bias toward posterior-to-anterior phase propagation. Colored bars represent regions of significant difference from baseline (q<0.001). Number of electrode pairs in each ROI pair shown. See also Supplementary Figure 3.

## Discussion

We have identified a visually-induced, macroscale, spatiotemporal propagation of phase modulation from posterior-to-anterior in human ventral occipitotemporal cortex. Strong phase modulation is seen almost ubiquitously across ventral occipitotemporal cortex in both hemispheres; from occipital pole to anterior fusiform. Following a visual event each region within the visual pathway displays a predictable, stereotyped phase progression. These findings were highly repeatable across a diverse range of visual stimuli with varying task demands and attentional loads and in only partially overlapping patient populations. We observed an approximate Euclidean propagation velocity of 0.8-1.0 m/s. This is comparable to the conduction velocities of unmyelinated horizontal projection axons in macaques (0.1-0.6 m/s) (Davis et al., 2020) and is much slower than would be expected from monosynaptic white matter transmission (>15 m/s) (Ferraina et al., 2002). This further implies that this phase propagation likely occurs via sequential cortico-cortical connections (Muller et al., 2014).

Previous work has shown the organization of spontaneous oscillations as large scale travelling waves (Bahramisharif et al., 2013; Muller et al., 2016; Zhang et al., 2018) and small scale travelling waves within early visual cortex (Muller et al., 2014; Sato et al., 2012). We expand on this work, characterizing a visually-evoked, macroscale phase propagation across the entire ventral visual pathway that likely indexes feedforward communication (Bastos et al., 2015), long-range cortico-cortical connections (Muller et al., 2014) and integration of information over broad cortical areas (Sato et al., 2012; Zhang et al., 2018).

A phase reset following visual input has been demonstrated in human (Mormann et al., 2005; Rizzuto et al., 2003; Tesche and Karhu, 2000) and macaque (Jutras et al., 2013) hippocampi. It is possible that this is also initiated as part of this sequence of ventral phase modulations. During visual exploration this hippocampal phase reset consistently occurs following saccadic eye movements (Jutras et al., 2013) suggesting that ventral phase modulation may occur following each saccade during naturalistic viewing.

Extracranial recordings of study phase modulation are an aggregate of many sources, limiting insight into underlying generators. Further, the relative inaccessibility of electrical fields generated by the ventral temporal surface (Goldenholz et al., 2009) has limited studies of visual phase modulation to early visual cortex. This has precluded insight into the spatiotemporal topography of visually-induced phase modulation in the human ventral visual pathway. Our results are supportive of the phase reset model of event related potential (ERP) generation (Klimesch et al., 2007; Makeig et al., 2002; Sauseng et al., 2007; Yeung et al., 2004). According to the phase reset model, the reset of instantaneous phase of ongoing oscillations, tending toward a fixed value upon presentation of a stimulus, results in the ERP by creating a coherent superposition of phase components (Iemi et al., 2019). Simulations of the underlying generators of the ERP suggest that early components (<200 ms), corresponding to the time course of ITC seen here, primarily represent feed-forward influences, with later ERP components being influenced by top-down modulation (Garrido et al., 2007).

## Conclusion

We demonstrate the existence of visually-induced macroscale posterior-to-anterior propagation of phase modulation across the human ventral occipitotemporal cortex. We propose this represents a feed-forward spread of information via horizontal cortico-cortical connections throughout the ventral visual pathway.

## Materials and Methods

### Participants

82 patients (44 male, 18-60 years, 16 left-handed) undergoing implantation with intracranial electrodes for seizure localization of pharmaco-resistant epilepsy participated after giving written informed consent. Participants with significant additional neurological history (e.g. previous resections, MR imaging abnormalities such as malformations or hypoplasia) were excluded. All experimental procedures were reviewed and approved by the Committee for the Protection of Human Subjects (CPHS) of the University of Texas Health Science Center at Houston as Protocol Number HSC-MS-06-0385.

### Electrode Implantation and Data Recording

Data were acquired using either subdural grid electrodes (SDEs; 14 patients) or stereotactically placed depth electrodes (sEEGs; 68 patients). SDEs were subdural platinum-iridium electrodes embedded in a silicone elastomer sheet (PMT Corporation; top-hat design; 3mm diameter cortical contact), surgically implanted via a craniotomy following previously described methods (Pieters et al., 2013; Tandon, 2012; Tong et al., 2020). sEEGs were implanted using a Robotic Surgical Assistant (ROSA; Medtech, Montpellier, France) (Rollo et al., 2020; Tandon et al., 2019). Each sEEG probe (PMT corporation, Chanhassen, Minnesota) is 0.8 mm in diameter, with 8-16 electrode contacts, each contact being a 2mm long platinum-iridium cylinder separated from the adjacent contact by 1.5 − 2.43 mm. Each patient had 12-20 such probes implanted. Following surgical implantation, electrodes were localized by co-registration of pre-operative anatomical 3T MRI and post-operative CT scans in AFNI (Cox, 1996). Electrode positions were projected onto a cortical surface model generated in FreeSurfer (Dale et al., 1999), and displayed on the cortical surface model for visualization (Pieters et al., 2013). Intracranial data were collected during research experiments starting on the first day after electrode implantation for sEEGs and two days after implantation for SDEs. Data were digitized at 2 kHz using the NeuroPort recording system (Blackrock Microsystems, Salt Lake City, Utah), imported into Matlab, and visually inspected for line noise, artifacts and epileptic activity. Of a total of 11,164 electrodes, 3,919 electrodes with obvious line noise or localized in proximity to sites of seizure onset were excluded. Each electrode was re-referenced to the common average of the remaining channels. Trials contaminated by interictal epileptic spikes were discarded.

### Signal Analysis

An ROI encompassing the entire occipital lobe and the ventral temporal surface, but excluding parahippocampal and entorhinal regions, was then applied, restricting us to 1,929 electrodes which form the basis of this work. These electrodes were grouped into sub-ROIs within 20 mm intervals along the y-axis of Talairach space, from −100 mm to 20 mm (Figure 1D-F).

Raw data was notch filtered to remove line noise (zero-phase 2nd order Butterworth band-stop filters). Phase information was extracted from the down-sampled (200 Hz) and wide band-pass filtered data (2 – 30 Hz; zero-phase 8th order Butterworth band-pass filter) using the ‘generalized phase’ method (Davis et al., 2020) with a single-sided Fourier transform approach (Marple, 1999). This method captures the phase of the predominant fluctuations in the wideband signal and minimizes filter-related distortion of the waveform.

Inter-trial Phase Coherence (ITC) was calculated as the absolute vector length of the mean of unit vectors with the instantaneous phase of stimulus-aligned trials (Figure 1G,H). Phase locking value (PLV) was calculated identically to ITC but instead using phase difference between two electrodes. ITC and PLV were computed as the median value of 50 iterations of a random 60 trials. ITC and PLV were baselined −1,000 to −100 ms before each stimulus. PLV was calculated between pairs of electrodes in separate ROIs, within patient, within hemisphere. PLV statistics were performed using a bootstrapped null distribution, randomly re-pairing trials between electrodes, using 5,000 repetitions. Phase probability distributions were quantified, using 30 equally spaced bins between − π and π, and mutual information (MI) was calculated based on these distributions at each time point, baselined −1,000 to −100ms before each stimulus. Asymmetry index (AI) was calculated based on the lagged PLV results at each time point as *AI* = *log*_*2*_(*PLV*_*pa*_/*PLV*_*ap*_), where PLV_pa_ represents the mean PLV between electrode *p* at time *t* and electrode *a* between time t and time *t* + 200 ms.

### Experimental Design and Statistical Analysis

Patients read single words (n = 40 patients) or identified celebrity faces (n = 57 patients). Stimuli were presented on a 2,880 × 1,800, 15.4” LCD screen positioned at eye-level, 2-3’ from the patient. Additional details of each experiment are below:

#### Word Reading

Participants read aloud monosyllabic words (e.g. dream) and pseudowords (e.g. meech) presented in lower-case Arial font, 150 pixels high (∼2.2° visual angle) (Figure 1A) (Woolnough et al., 2021a). Stimuli were presented using Psychophysics Toolbox (Kleiner et al., 2007) run in Matlab. Each stimulus was displayed for 1,500 ms with an inter-stimulus interval of 2,000 ms. Stimuli were presented in two recording sessions, each with 160 stimuli in a pseudorandom order with no repeats.

#### Face Recognition

Participants were presented with greyscale photos of famous faces and asked to name them aloud (Figure 1B) (Kadipasaoglu et al., 2017; Woolnough et al., 2020). Stimuli were matched for luminance and contrast and presented, using Python 2.7, at a size of 500 × 500 pixels (∼7.5° visual angle) for 2,000 ms, with an inter-stimulus interval of 6,000 ms. Stimuli were presented in one recording session, with 60-125 faces presented in pseudorandom order with no repeats. Participants who correctly identified less than 20 stimuli were excluded.

Linear regressions were performed using robust regression models (Matlab function *fitlm* using *‘RobustOpts’*; iteratively reweighted least squares with bisquare weighting) (Welsch, 1977), to minimize the effect of outliers and to make fewer assumptions than traditional least squares linear regression models. To calculate onset latencies, we determined the first time point per electrode, within 600 ms of stimulus onset, where the first derivative of the ITC exceeded 3.5 standard deviations (∼p<0.001) from baseline. We then used a regression model, predicting onset latency based on position in the y (anteroposterior) axis, derived from the surface-based co-registration. All temporal analyses were corrected for multiple comparisons using a Benjamini-Hochberg false-detection rate (FDR) threshold of q<0.001.

## Acknowledgements

We express our gratitude to all the patients who participated in this study; the neurologists at the Texas Comprehensive Epilepsy Program who participated in the care of these patients; and the nurses and technicians in the Epilepsy Monitoring Unit at Memorial Hermann Hospital who helped make this research possible. This work was supported by the National Institute for Deafness and other Communication Disorders DC014589 and National Institute of Neurological Disorders and Stroke NS098981.

## Author Contributions

Conceptualization: OW, NT; Methodology: OW, NT; Data curation: OW, PSR, ZR; Software: OW; Formal Analysis and Visualization: OW; Writing – Original Draft: OW; Writing – Review and Editing: OW, KJF, NT; Funding Acquisition: NT; Neurosurgical Procedures: NT.

## Supplementary Figures

**Supplementary Figure 1:**
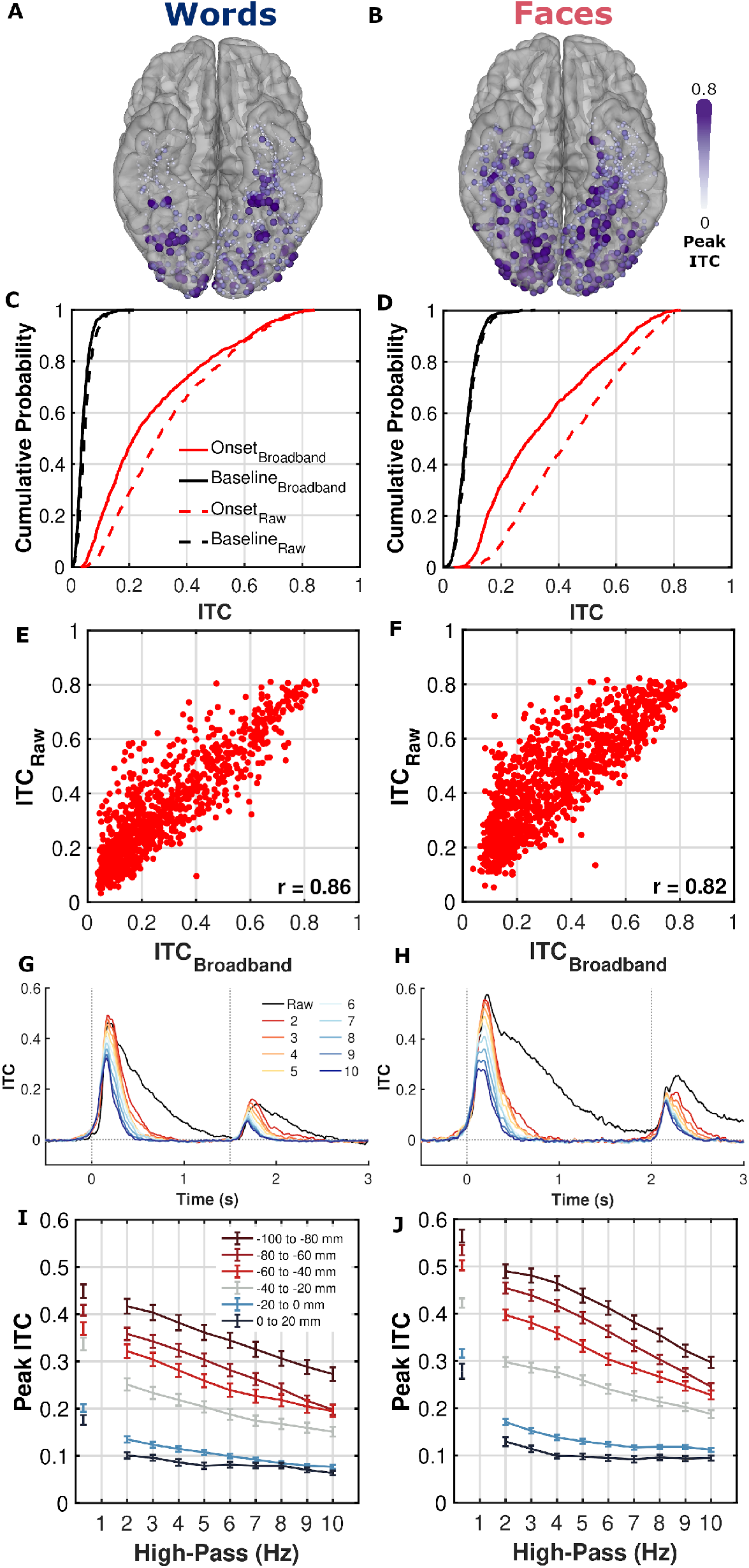
Comparison of Broadband Low Frequency and Raw Signal Analyses. (A,B) Spatial distribution of peak ITC for each electrode in the raw (0.3 – 100 Hz) signal. (C,D) Cumulative probability distribution of raw and broadband ITC. (E,F) Relationship of raw and broadband ITC. (G,H) Temporal ITC profile with varying high pass filter parameters. Same electrodes as Figure 3D,E. (I,J) Changes in peak ITC within each spatial ROI for the raw signal (0.3 Hz high pass) and varying high pass filtered signals (2 - 10Hz). Analyses shown for the words (A,C,E,G,I) and faces (B,D,F,H,J) tasks.

**Supplementary Figure 2:**
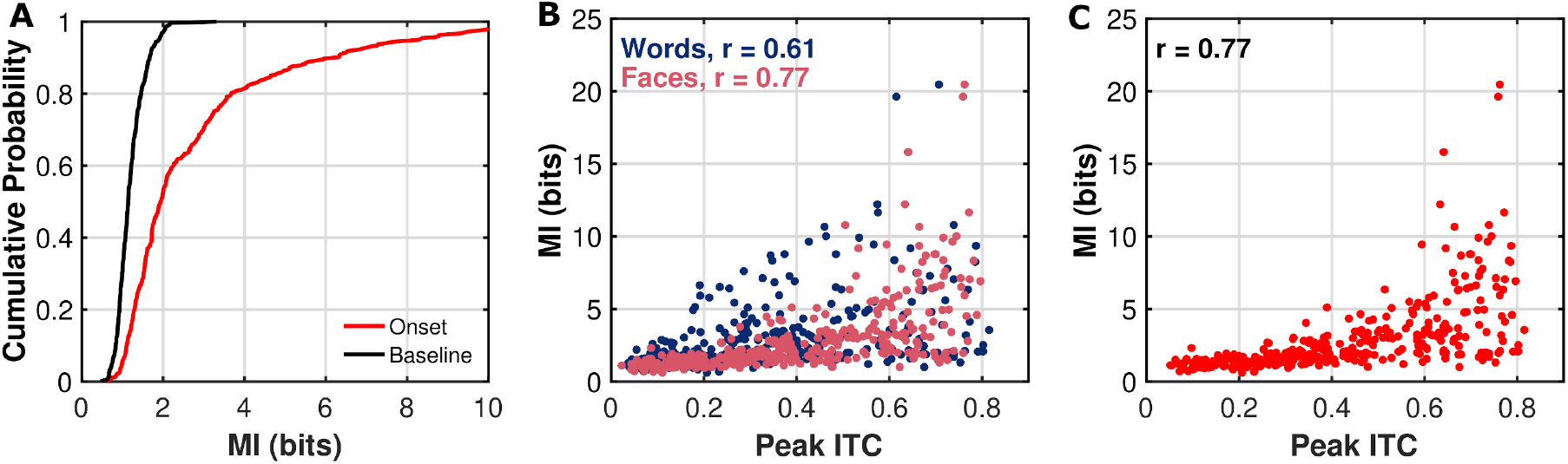
Distribution of Mutual Information (MI). (A) Cumulative distribution function of peak MI within the onset window (0 to 600 ms; red) or during the baseline window (−500 to −100 ms; black). (B,C) Relationship between peak MI and (B) peak ITC in the words and faces tasks or (C) peak ITC across both tasks, within individual electrodes.

**Supplementary Figure 3:**
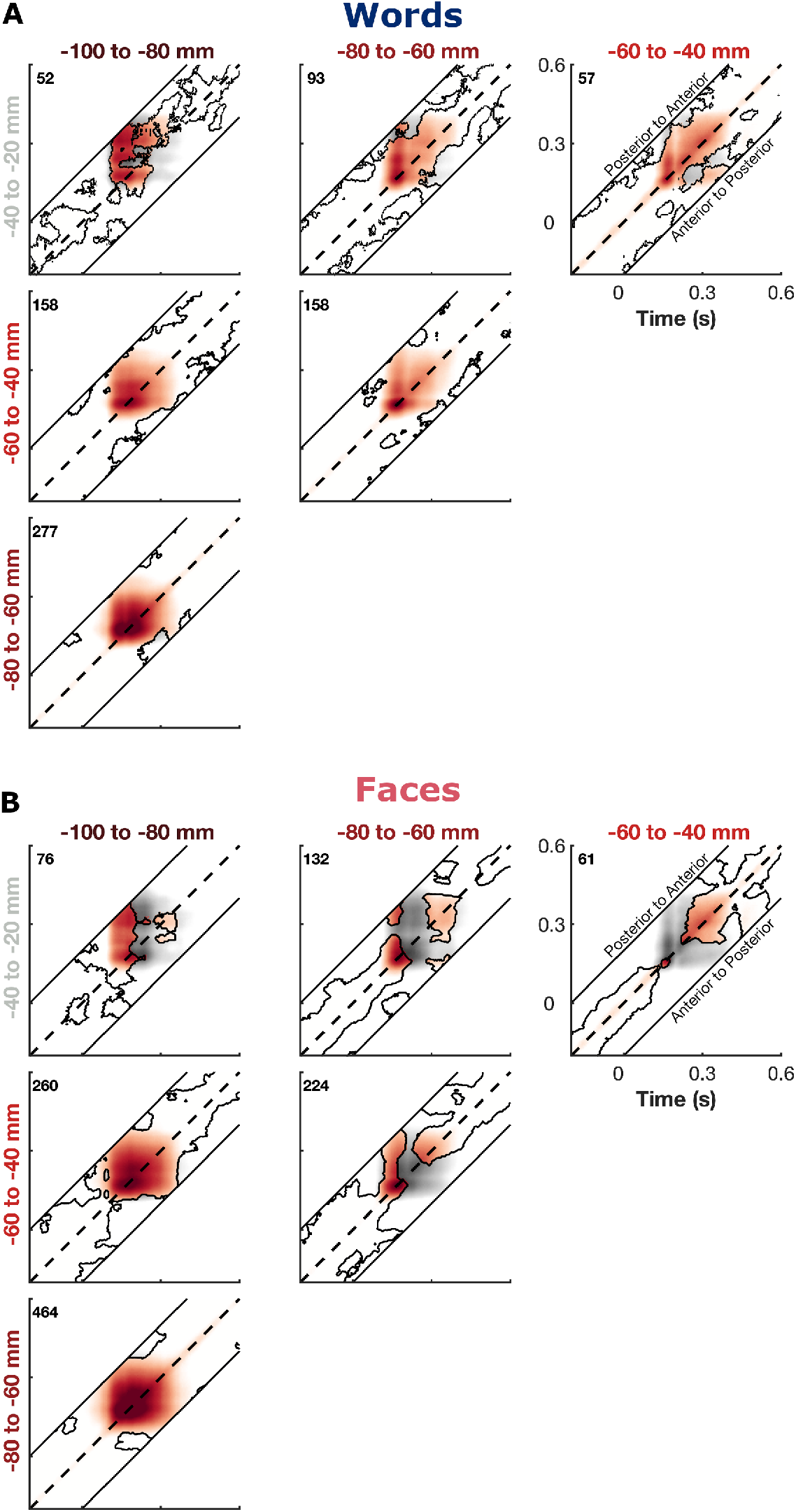
Lagged Phase Locking Value. PLV was calculated between the instantaneous phase of the posterior electrodes (columns) and the time lagged phase of the anterior electrodes (rows; −200 to 200 ms) for the words (A) and faces (B) tasks. Greater PLV above the center diagonal (dashed line) represents a greater ability to predict future phase of the anterior electrodes based on the current phase of the posterior electrodes. Areas of significant PLV (q< 0.05) above a trial shuffled null distribution are indicated by contours and non-significant areas are desaturated. Number of electrode pairs in each ROI pair shown.

